# Disruptions in effort-based decision-making following acute optogenetic stimulation of ventral tegmental area dopamine cells

**DOI:** 10.1101/2020.12.09.417832

**Authors:** Benjamin R. Fry, Nathan T. Pence, Andrew McLocklin, Alexander W Johnson

## Abstract

The dopamine system has been implicated in decision-making particularly when associated with effortful behavior. We examined acute optogenetic stimulation of dopamine cells in the ventral tegmental area (VTA) as mice engaged in an effort-based decision-making task. Tyrosine hydroxylase-Cre mice were injected with Cre-dependent ChR2 or eYFP control in VTA. While eYFP control mice showed effortful discounting, stimulation of dopamine cells in ChR2 mice disrupted effort-based decision-making by reducing choice towards the lever associated with a preferred outcome and greater effort. Surprisingly, disruptions in effortful discounting were observed in subsequent test sessions conducted in the absence of optogenetic stimulation, however during these sessions ChR2 mice displayed enhanced high choice responding across trial blocks. These findings suggest increases in VTA dopamine cell activity can disrupt effort-based decision-making in distinct ways dependent on the timing of optogenetic stimulation.

---

The neurotransmitter dopamine has been implicated in a wide-range of learning and motivational processes including effort-based decision-making. At present, many studies investigating dopamine’s role in this form of decision-making have found a dichotomous pattern of results in which receptor antagonism (Robles & Johnson, 2017; Bryce & Floresco, 2019) or depletion (Cousins & Salamone, 1994; Mingote et al., 2005) of dopamine results in a biasing of behaviors away from more preferred rewards associated with increased effort. Conversely, dopamine stimulation (Bardgett et al., 2009; Wardle et al., 2011) or transgenic overexpression of dopamine receptors (Trifilieff et al., 2013) tends to invigorate responding for outcomes associated with higher effort. While much has been learned, a limiting factor with these approaches is that they lack temporal specificity, thus constraining resultant interpretations. We employed optogenetics to determine the effect of acute ventral tegmental area (VTA) dopamine stimulation at the timepoint of decision-making. Such an approach allows for a focus strictly on the role of dopamine in evaluating choices with different cost-benefit outcomes, without the potentially confounding factor of consistently altered dopamine signaling throughout the decision-making test.

Twenty-three mice expressing Cre recombinase under the control of the tyrosine-hydroxylase promoter were produced via outbreeding with wild-type C57BLJ mice (Jackson Laboratory, Bar Harbor, ME) under the auspices of the Michigan State University Institutional Animal Care and Use Committee. Mice were housed by sex, up to five per cage pre-surgery. At ten weeks of age, mice underwent stereotaxic surgery in which 0.5 μl of either AAV5-Ef1α-DIO-ChR2-eYFP (n = 10♂, n = 8♀) or AAV5-Ef1α-DIO-eYFP (n = 5♂, n = 3♀; Vector Biolabs, Malvern, PA) was virally infused unilaterally at the level of the VTA (AP −3.08, ML +/−0.6, DV - 4.5 mm) in a manner counter-balanced for hemisphere. Optic fiber ferrule tips (200μl core, 4.1mm; Thorlabs, Newton, NJ) were implanted dorsal (≈ 0.3 mm) to the injection site and affixed with dental acrylic (Lang Dental Manufacturing Co, Wheeling, IL). Mice were given four weeks post-surgery to recover and to allow for sufficient viral transfection, during which time they were singly housed, which continued for the duration of the experiment in order to avoid ferrule tip damage.

Following recovery, mice were food deprived to 90% of their free-feeding weight by limiting access to a single daily portion of lab chow. Subsequently, mice were trained in operant chambers (Med Associates, St Albans, VT) where in separate sessions, responses on one lever led to 50 μl delivery of a preferred high value reinforcer (e.g., left lever→orange-flavored 20% sucrose), whereas a different lever resulted in delivery of the lower value outcome (e.g., right lever→grape-flavored 5% sucrose). Apart from reversal testing, these lever contingencies remained fixed throughout and were counterbalanced across viral groups. Training continued until mice hit a criterion of 25 responses for each lever in a single session. Next, mice were trained on the effort-based decision-making task at which time they were tethered, however no optical stimulation occurred. These approximately 1hr sessions were divided into four blocks consisting of ten trials each and were conducted once per day for 10-12 days. Within each block, mice first received four forced trials (two low effort, two high effort, randomly ordered) in which only one lever was extended. Subsequently, mice received six choice trials where both levers extended simultaneously. Across blocks, a single press on the low effort, low choice lever led to the delivery of the low value reinforcer (i.e., fixed-ratio 1; FR-1). Alternatively, responses on the high choice lever led to the delivery of the high value reinforcer, which gradually required more responses to acquire across blocks: block 1 = FR-1; block 2 = FR-5; block 3 = FR-20; block 4 = FR-40. For choice trials, the first response during block 1 trials led to the retraction of both levers, whereas during blocks 2-4 the first response on the high choice lever led to the retraction of the low choice lever. Each trial and block were separated by a fixed 60 s interval, and during the trials if mice failed to perform a lever response across a 60 s interval the trial was omitted.

Following training, mice underwent two optogenetic testing days. During these stimulation sessions, light intensity was initially calibrated to emit ≈ 20mW from the tip of the optical fiber, with 1 s optogenetic stimulation of VTA dopamine cells (473 nm, 5 ms pulses at 20Hz) beginning 5 s prior to each choice trial, which was delivered via a waveform generator (Agilent Technologies, Santa Clara, CA) integrated into the Med Associates apparatus. Subsequently, half the mice from each viral condition received one post-stimulation test, whereas the other half received two post-stimulation tests. Finally, in order to control for any potentially confounding effects of dopamine stimulation on contingency learning, a small cohort of mice (n = 4) received additional training wherein response-outcome contingencies were reversed (i.e. low effort lever always yielded 20% sucrose under FR1, while high effort lever yielded 5% sucrose under increasing effortful action). Following ten sessions of training, these mice received optogenetic testing as described above, albeit with the new response-outcome contingencies.

Upon conclusion of testing, mice received an intraperitoneal injection of sodium pentobarbital (100 mg/kg). Subsequently, they were perfused transcardially with 0.9% saline followed by 4% cold paraformaldehyde (Sigma-Aldrich, St. Louis, MO) in 0.1 M phosphate buffer (PB). Brains were extracted and post-fixed in a 12% sucrose, 4% paraformaldehyde solution, for ≈ 24 hr at 4°C. Brains were then extracted, sliced at 30 μm using a freezing microtome and stained using a mouse-anti-tyrosine-hydroxylase primary (Catalog# MAB318; MilliporeSigma) in addition to a rabbit-anti-GFP primary (1:1000; Catalog# MAB318; MilliporeSigma) with donkey-anti-rabbit-488 (1:500; Catalog# A21206; Invitrogen) and donkey-anti-mouse-568 (1:500; Catalog# A10037; Invitrogen) corresponding to ChR2 and TH positive cells, respectively. Imaging of the VTA (Fig. 1a) was carried out using a Nikon A1 laser-scanning confocal microscope (Nikon Instruments Inc, Melville, NY). Viral targeting and co-localization were scored qualitatively using separate raters (Fig. 1b). At this time, n = 2 ♂ and n = 2♀ from group ChR2 were excluded from analysis due to poor targeting.

**Figure 1.**
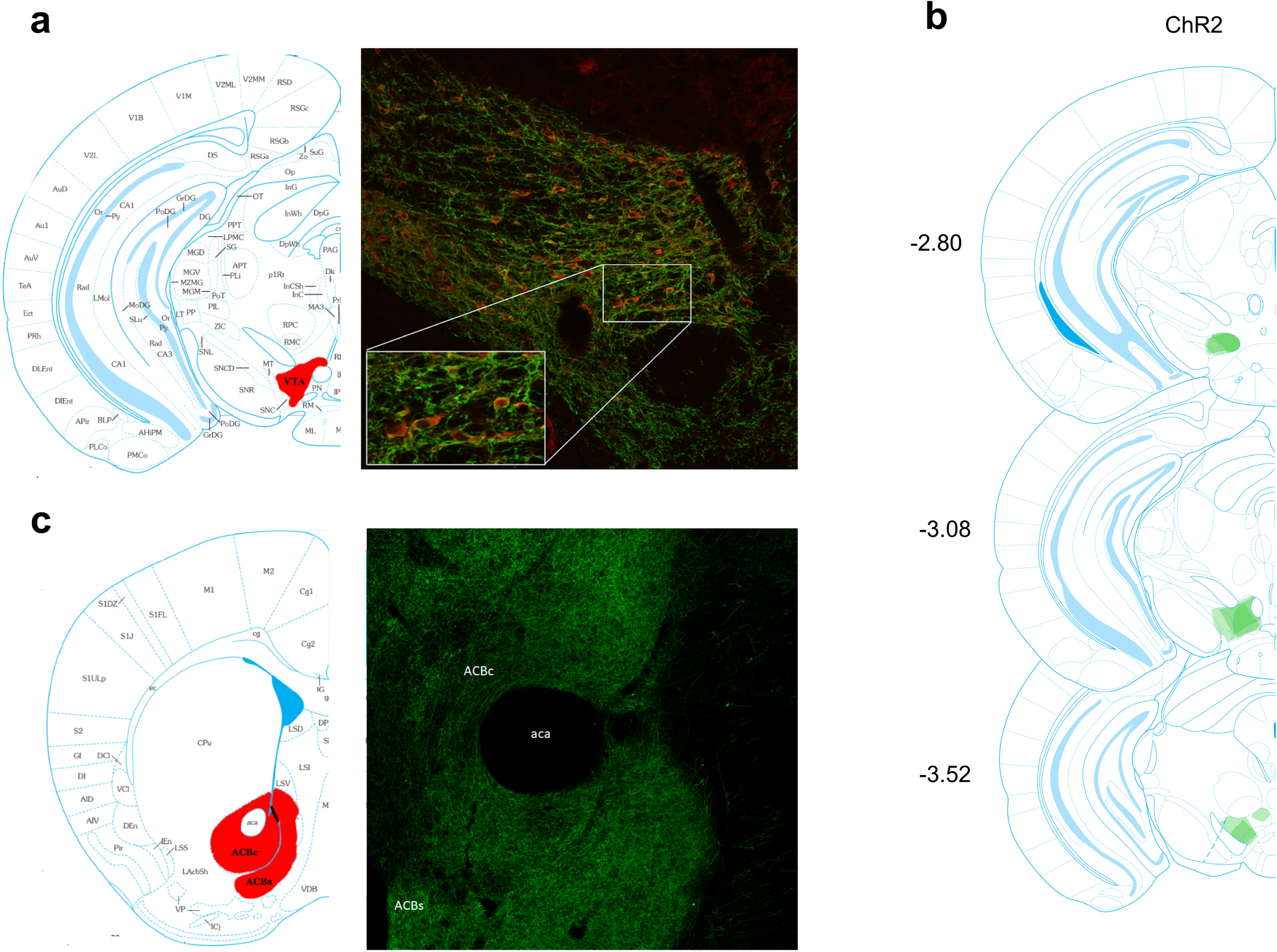
Representative photomicrograph showing immunohistochemical. **(a)** verification of Cre-dependent ChR2 (green) in tyrosine-hydroxylase positive cells (red) in VTA. **(b) Quantified eYFP expression of ChR2 in VTA is displayed where light shading represents minimal spread and darker shading represents maximal spread of viral expression at each level. (c)** Dense viral transfection was noted in terminals in nucleus accumbens. ACBs = nucleus accumbens shell; ACBc = nucleus accumbens core; aca = anterior commissure.

For data analysis, for each trial block the percentage choice on the high choice lever with omissions excluded was averaged for the final two training sessions (pre-stimulation), the optogenetic test sessions (stimulation) and the proceeding non-stimulated session(s) (post-stimulation). These data were subjected to a three-way repeated measures ANOVA with a between-subjects variable of virus (ChR2 vs eYFP) and within-subjects variable of session type (pre-stimulation, stimulation, post-stimulation) and block (1-4). Follow-up virus X block interactions were carried out for each session separately, with tests of simple main effects used to determine the nature of any significant interactions. Post-hoc Bonferroni correction to control for multiple comparisons was used to determine significant main effects of either block or session. Finally, due to a lack of variance in a number of the test blocks and non-uniform distribution, analysis of the omission and contingency-reversal data was conducted using Wilcoxon matched pairs test. The α level for significance was .05 and all analyses were conducted using Statistica (Statsoft, Tulsa, OK).

The data of primary interest are depicted in Figure 2. For the choice test data (Fig’s 2a-c), analyses revealed a significant virus X session X block interaction (F(6,120) = 2.38, p<0.05). Subsequent virus X block ANOVA for the pre-stimulation session (Fig. 2a) revealed a main effect of block only (F(3,60) = 3.58, p=0.01) due to significant discounting of the high effort reward in all mice on block 4 relative to blocks 1 and 2 (p’s≤0.02). By comparison, optogenetic stimulation of DA cells significantly disrupted choice test responding (Fig. 2b). ANOVA revealed a main effect of virus (F(1,20) = 5.12, p<0.05) and block (F(3,60) = 3.55, p=0.01). Planned comparisons revealed a significant reduction on high choice responses in ChR2 mice relative to eYFP controls on block 1 (F(1,20) = 7.51, p=0.01) and a tendency for group differences on blocks 3 and 4 (smallest F-value; block 4, F(1,20) = 3.28, p=0.08). This difference in performance between the groups could not be attributed to non-specific effects of laser stimulation on motoric action as omissions (Fig. 2e) were comparable across trial blocks (largest Z-value; block 2, *Z* = 0.54, p=0.58). Strikingly, in the sessions proceeding optogenetic testing, ChR2 mice displayed disruptions in effortful discounting as they continued performing on the high choice lever irrespective of increases in the effort required to obtain reward (Fig. 2c). ANOVA revealed a significant block X virus interaction (F(3,60) = 4.66, p=0.005) due in part to significant reduction in high choice responses on block 4 in eYFP relative to ChR2 mice (F(1,20) = 4.72, p<0.05). Moreover, eYFP mice discounted the 20% sucrose reinforcer as the amount of effort increased to FR-40 relative to performance during all other trial blocks (smallest F-value; block 1 vs 4, F(1,20) = 4.54, p<0.05). By contrast, ChR2 mice displayed comparable responding for the high choice lever across all trial blocks (F’s<1). These effects also did not reflect generalized motoric disruptions as omissions across trial blocks were comparable in ChR2 and eYFP mice (Fig. 2f ; largest Z-value; block 2, *Z* = 0.98, p=0.32). Finally, as the effects of optogenetic stimulation in ChR2 animals were present in the first block (Fig. 2b) we wanted to confirm that laser activation in this group did not disrupt the capacity of mice to encode and/or retrieve lever contingencies (Fig. 3). Analyses revealed that irrespective of dopamine VTA stimulation, mice continued to bias their choice performance towards the low choice lever, which at this stage of testing led to delivery of the preferred higher value reinforcer (largest Z-value; block 3, *Z* = 1.60, p=0.1).

**Figure 2.**
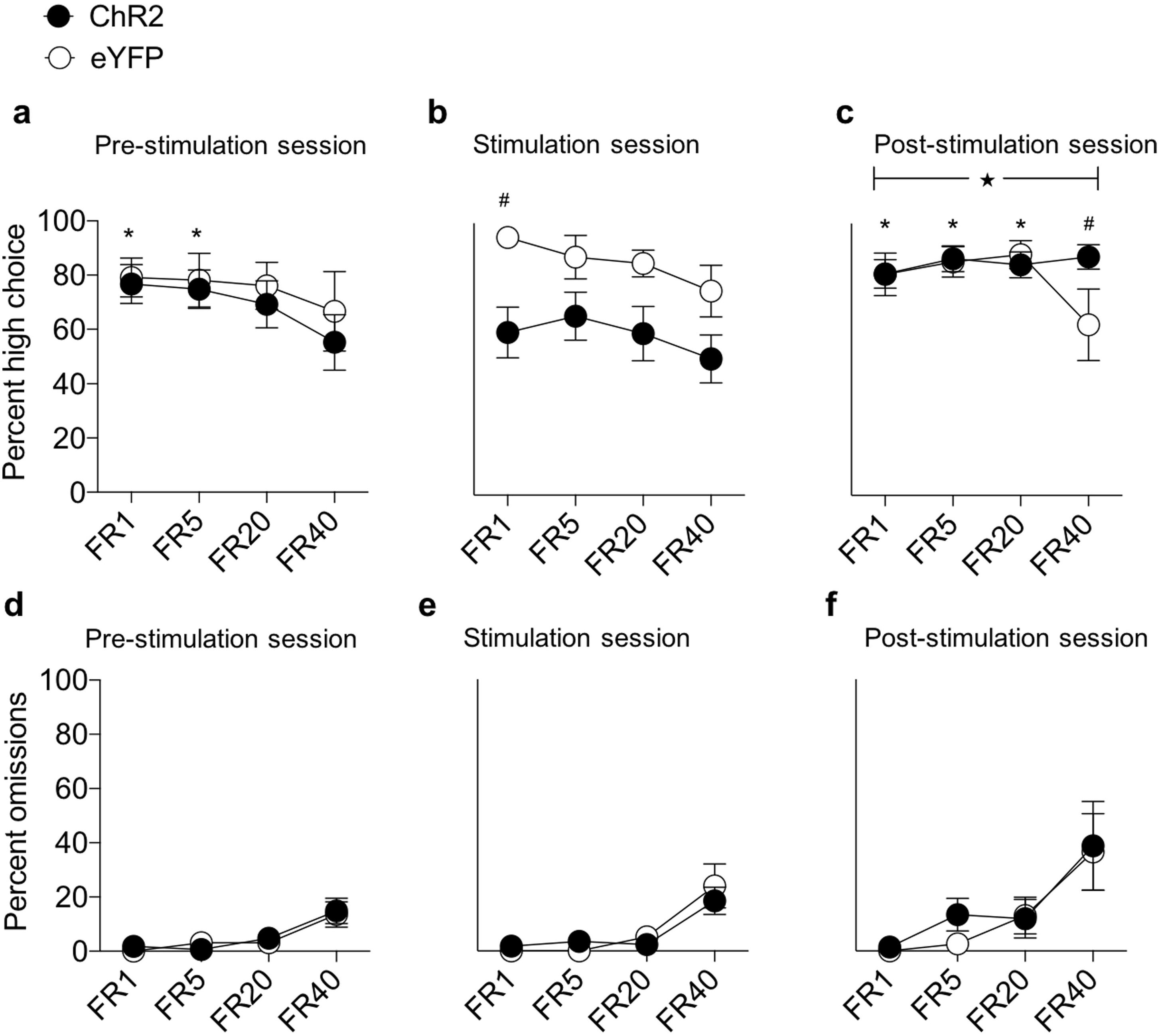
Optogenetic stimulation of VTA dopamine cells disrupts effort-based decision-making. **(a)** In the pre-stimulation decision-making test sessions prior to optogenetic stimulation, both ChR2 and eYFP mice displayed a comparable pattern of responding by reducing their high choice lever responses during the final block of FR-40 trials. Post-hoc Bonferroni contrasts revealed significant reduction in high choice lever responding in blocks 1 (# p<0.01) and 2 (# p=0.02) relative to block 4. **(b)** Optogenetic stimulation in ChR2 mice led to a reduced preference for the high choice lever across trial blocks. # indicates significant group difference during block 1 (p=0.01). **(c)** In the effort-based decision-making tests following optogenetic stimulation, post-session test choice performance reflected a disruption in effortful discounting in ChR2 mice as they persistently preferred the high choice lever across trial blocks. ★ indicates significant virus X block interaction (p=0.005), # indicated significant group differences during block 4. * indicates significant reduction in high choice responses for eYFP block 4 trials relative to blocks 1 (p=0.05), 2 (p=0.003) and 3 (p<0.001). **(d-f)** Omissions generally increased during FR-40 trials across **(d)** pre-stimulation, **(e)** stimulation and **(f)** post-stimulation test sessions, though importantly no group differences were noted (p’s>0.17).

**Figure 3.**
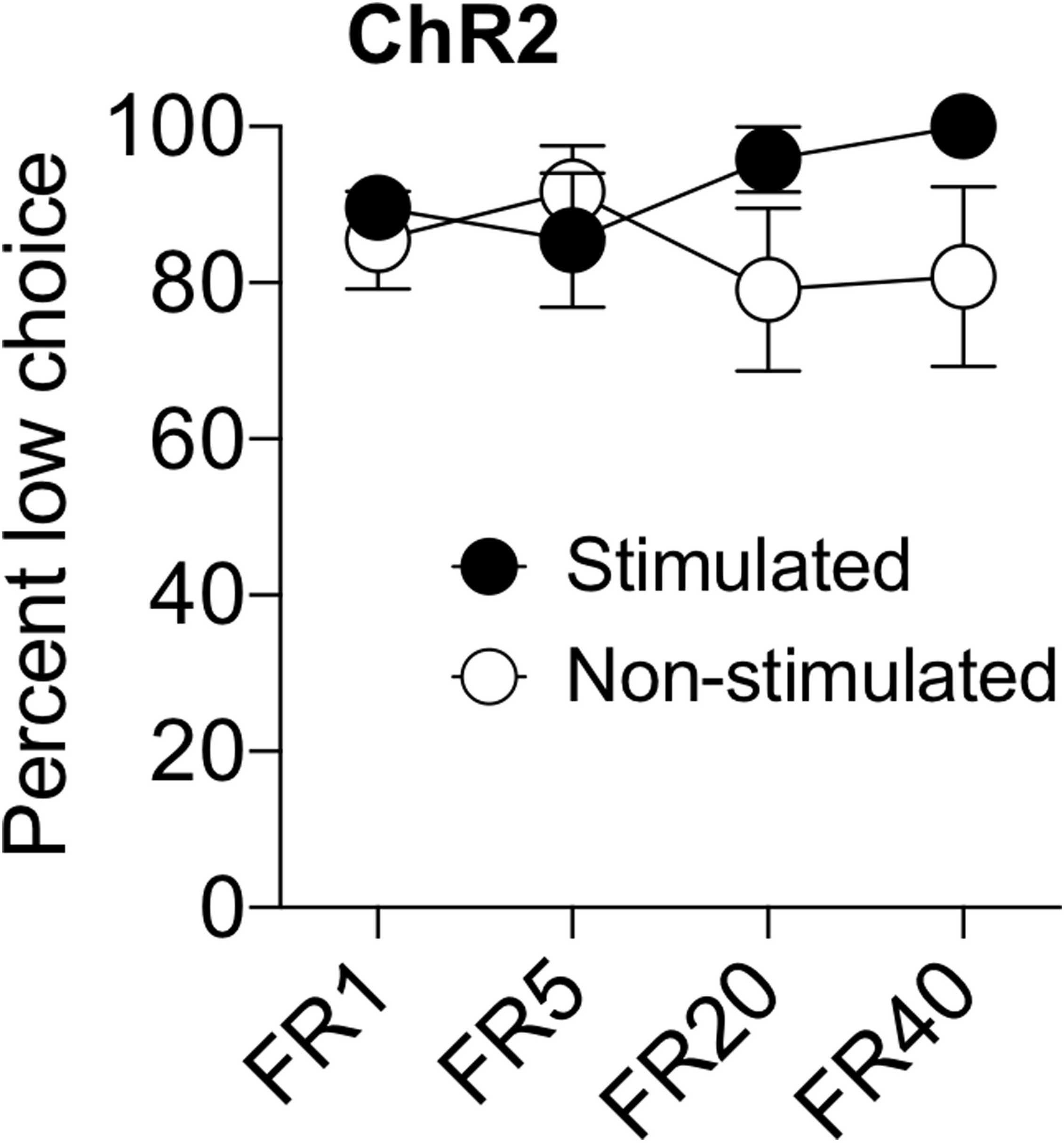
Following reversal of the lever contingencies such that the low effort lever always produced the preferred high value outcome, ChR2 mice continually maintained their preference for the low effort lever irrespective of laser stimulation or trial block.

Previous studies have shown that dopamine depletion or antagonism reduces performance following increases in the effort required to obtain reward (Robles & Johnson, 2017; Salamone et al., 1991; Bardgett et al., 2009; Nowend et al., 2001; Mingote et al., 2005), whereas facilitations in dopamine signaling bias performance towards effortful actions (Wardle et al., 2011; Bardgett et al., 2009; Trifilieff et al., 2013). These findings suggest dopamine plays an integral role in decision-making when options differ in their costs and are broadly consistent with the sensorimotor activational hypothesis (Salamone et al., 2007), which posits that dopamine promotes action generation and the prolongation of effortful behavior. However, a limitation of these studies is their reliance on perturbations of dopamine function over relatively protracted timeframes.

In the current study, when acute optogenetic stimulation of dopamine cells transiently preceded effort-based decision-making, mice displayed a general reduction in choice performance for the high effort lever. These findings contrast with the vast majority of decision-making studies that indicate increases in dopamine promote responding for more effortful actions. Nevertheless, several findings suggest nuances in this relationship. For instance, peripheral injections of amphetamine in rats significantly decreased lever responding for preferred food pellets (Cousins et al., 1994) and reduced preference for a high value food outcome as the amount of effort required to obtain it increased (Floresco et al., 2008). Similarly, administration of the D2/D3 receptor agonist quinpirole in rats (Depoortere et al., 1996), or transgenic overexpression of post-synaptic striatal dopamine D2 (Drew et al., 2007) or D3 (Simpson et al., 2014) receptors produce progressive ratio deficits. Moreover, D2 receptor overexpression also reduced the willingness to work for palatable food in a cost-benefit decision-making task (Filla et al., 2018). More targeted pharmacological approaches in rats likewise reveal a similar pattern of performance, whereby administration of a D2/D3 receptor agonist (but not D3 agonism alone) in the nucleus accumbens reduced choice responding for the high effort lever during effort-based decision-making (Bryce & Floresco, 2019). Given that VTA injections of ChR2 was trafficked to a number of targets including the nucleus accumbens (Fig. 1c), this potentially suggests mesostriatal modulation underlying the disruptions in effort-based decision-making (Fig. 2b). Notably, the influence of optogenetic stimulation on choice behavior persisted beyond optogenetic testing such that in the post-stimulation sessions (i.e., when the laser was not activated), ChR2 mice displayed a persistent preference for the high choice lever that was not dampened by increases in effort (Fig. 2c). This disruption in effortful discounting is similar to the preponderance of pharmacological and transgenic findings in which dopamine stimulation or receptor overexpression augments responding for higher effort outcomes (Bardgett et al., 2009; Wardle et al., 2011; Trifilieff et al., 2013).

Our findings raise important questions regarding the role and consequences of acute mesencephalic dopamine cell stimulation on effort-based decision-making. Unlike past studies, acute stimulation of VTA dopamine cell activity reduced preference for the high value outcome even when effort costs were equated (i.e., during FR-1 block trials; Fig. 2a). This suggests optogenetic stimulation may have disrupted the reinforcing efficacy of the higher value reinforcer, and/or the animals’ capacity to discriminate between lever contingencies. However, this interpretation is unlikely given that when mice were tested under conditions where the higher value outcome was continually paired with low effort, mice maintained their preference for the larger preferred reward across trial blocks independent of VTA stimulation (Fig. 3). In addition, findings from either the stimulation or post-stimulation tests are unlikely to reflect either baseline group differences in performance (Fig. 2a; supplemental Fig. 1), or gross changes in motoric output and motivation, as omissions were comparable across all test days (Fig. 2d-f). Beyond these more prosaic interpretations, it is worthwhile considering that in addition to its activational effects, dopamine acts a value-based prediction error signal critical for reward learning (Schultz, 1997; Waelti et al., 2001; Bayer & Glimcher, 2005; Eshel et al., 2016; Steinberg et ., 2013; Pessiglione et al., 2006; Bayer et al., 2019). Accordingly, in the optogenetic stimulation sessions decision-making performance might be influenced by disruptions in associative mechanisms controlled by reward prediction error signals. Furthermore, more recent studies suggest dopamine transients can be uncoupled from model-free reinforcement value signals to encode associative information that is computationally detailed in nature (e.g. its sensory features; Gardner et al., 2018; Sharpe et al., 2019). This in mind, the current findings also suggest that dopamine functions beyond a pure value-based signal, to include recall of past, and/or anticipation of detailed upcoming contingencies, which may bias performance towards actions that overall have the highest immediacy of reinforcer delivery. This could serve to direct actions towards the low effort lever where execution of a single lever press consistently results in immediate reward delivery even under cost-benefit conditions where larger rewards may be available. Alternatively, the pattern of responding during laser activation may be an artifact of optogenetic stimulation of VTA dopamine transients, whereby endogenous phasic dopamine release that supports effortful action when produced by extension of the levers may have been attenuated as a result of the preceding optogenetic stimulation (e.g., via an insufficient refractory period). Regarding post-stimulation performance, our findings suggest that prior optogenetic stimulation subsequently enhanced responding for more rewarding outcomes when mice were tested without acute dopamine cell activation. Perhaps in the absence of prior dopamine cell activation, ChR2 mice were in an attenuated reward state during the post-stimulation decision-making task and compensated for this by enhancing their reward seeking for the highest value reward present (i.e., 20% sucrose). Moreover, these findings should be considered when developing future experimental designs, given that the longevity of post-stimulation disruptions in effortful discounting is currently unknown.

While our study provides important insight into the role of VTA dopamine transients on effort-based decision-making, a number of caveats should be acknowledged. As suggested above, this study would benefit from optical stimulation at different time points during the choice test. In addition, VTA dopamine efferents target numerous striatal (nucleus accumbens core and shell), limbic (basolateral amygdala) and prefrontal (orbitofrontal cortex, anterior cingulate cortex) sites that are implicated in decision-making (Cousins et al., 1996; Winstanley et al., 2004; Floresco & Ghods-Sharifi, 2007; Hauber & Sommer, 2009; Walton et al., 2009). Thus, photostimulation of axonal terminals in these downstream targets will provide additional insight into the circuit level effects of dopamine stimulation on effortful discounting. It should also be recognized that our use of TH-Cre mice to control dopamine cell function is somewhat confounded by expression of TH transgene in non-dopamine cells within the mesencephalon (Lammel et al., 2015; Stuber et al., 2015), consequently future approaches should employ DAT-Cre lines where ectopic expression is minimal. Finally, a number of relevant measures including latencies for deliberation time were not recorded.

Effort-based decision-making impairments have been noted in numerous settings including in patients with depression, schizophrenia and autism (Treadway et al., 2012, 2015; Damiano et al., 2012; Green et al., 2015), along with obese individuals (Mata et al., 2017). Although reduced dopamine transmission has been generally attributed to underlie these disruptions, results from our study add to a smaller yet important body of work indicating that aberrant elevations in dopamine signaling may impair cost/benefit decision-making. They are also consistent with contemporary accounts that indicate a more complex computational role for dopamine transients, which may include promoting action selection towards more immediate reward delivery. Surprisingly, the acute effects of dopamine stimulation endure beyond the stimulation test sessions by altering decision-making performance and invigorating reward-seeking for higher value and effortful rewards.

