## Supplemental text for "Disruptions in effort-based decision-making following acute optogenetic stimulation of ventral tegmental area dopamine cells"

**Supplemental Figure 1**. Across training mice gradually increased their preference for the high choice lever, and towards the end of training displayed a pattern of responding in which high choice lever responses gradually decreased as the effort schedule increased. Virus (eYFP, ChR2) X session (1-10) x block (1-4) ANOVA revealed a main effect of session (F(4,80) = p<0.001) and block (F(3,60) = 12.87, p<0.001). No effect of virus or interaction between the variables was noted (Largest F-value; session X block, F(12,240) = 1.31, p=.02).
