## Supplementary figures and images for "Disruptions in effort-based decision-making following acute optogenetic stimulation of ventral tegmental area dopamine cells"

### Supplemental figure

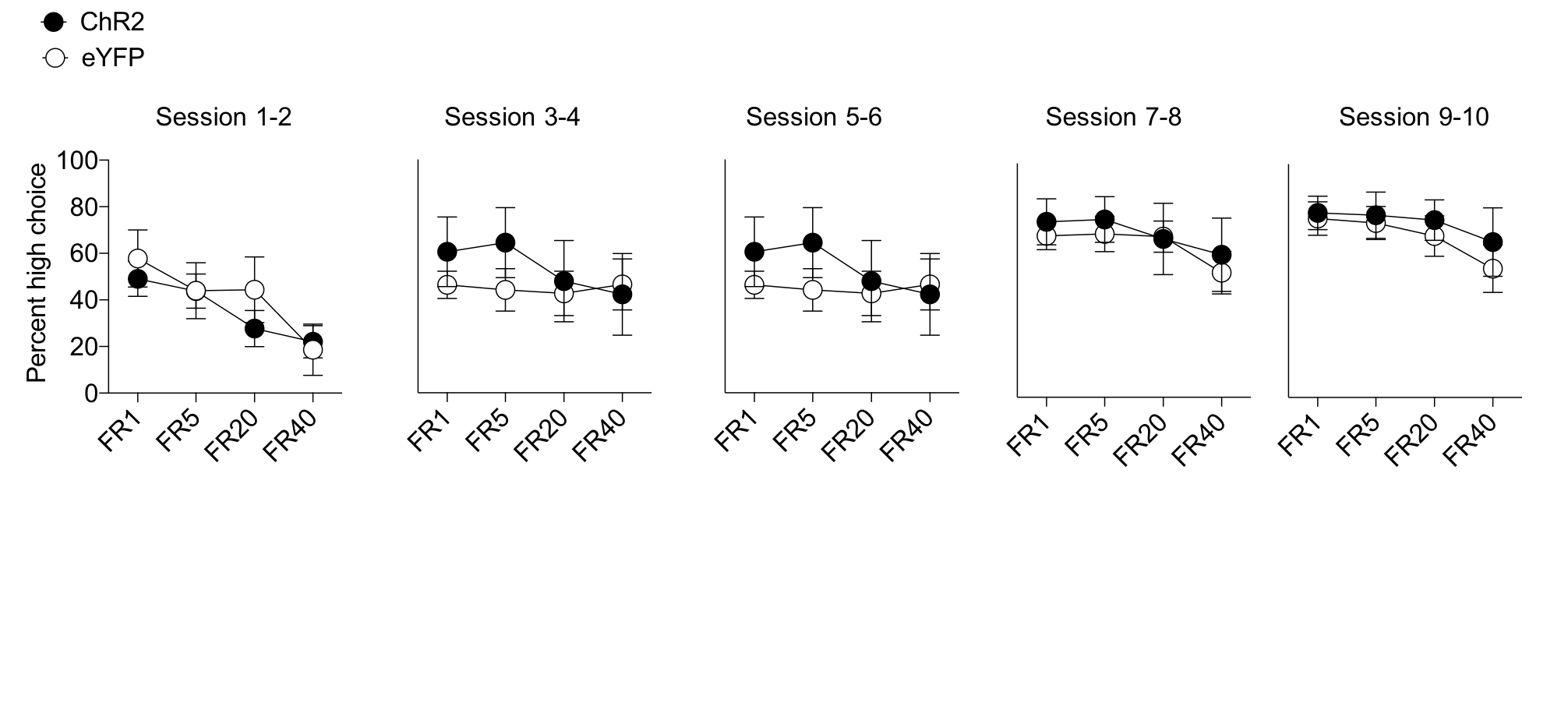
